# Benzyl isothiocyanate induces cardiomyocyte proliferation and heart regeneration

**DOI:** 10.1101/2021.09.08.459197

**Authors:** Akane Sakaguchi, Miwa Kawasaki, Yuichi Saito, Kozue Murata, Hidetoshi Masumoto, Wataru Kimura

## Abstract

Cardiomyocyte proliferation is an evolutionarily conserved mechanism that supports cardiac regeneration in vertebrates. Mammalian cardiomyocytes are arrested from the cell cycle shortly after birth, and therefore mammals lose the ability to regenerate injured myocardium for the rest of their lives. Pharmacological induction of cardiomyocyte proliferation has gained a lot of interest in recent years, as researchers strive to achieve heart tissue regeneration. Here we show that a small chemical, benzyl isothiocyanate (BITC), induced cardiomyocyte proliferation through activation of the cyclin-dependent kinase (CDK) pathway. BITC treatment also allowed heart regeneration in the infarcted neonatal heart, even after the regeneration period in mice. Furthermore, administration of BITC to adult mice in parallel with mild hypoxia (10% O_2_) induced cell cycle reentry and tissue regeneration in the adult heart. Our findings thus suggest that pharmacological activation of the CDK pathway using BITC, concurrently with the activation of hypoxia-related signaling pathways, may be a promising approach to inducing cardiac regeneration in patients with heart disease.

## Introduction

Mammalian cardiomyocytes have the ability to proliferate from the embryonic stage until the early neonatal stage, with most of them being arrested from the cell cycle shortly after birth^1^. While it has been shown that cardiomyocyte proliferation takes place in the adult mammalian heart, the rate of cell division is rare, only allowing the replacement of cardiomyocytes at a rate of less than 1% annually^2-4^. Therefore, adult mammalian heart cannot repair myocardial injury, resulting in the fact that ischemic heart disease is the leading cause of death in the world. As such, enhancement of cell cycle reentry for endogenous cardiomyocytes is of great interest for the replacement of the injured fibrotic scar tissue with healthy myocardium^5^. Activation of cardiomyocyte cell cycle via gene knockout^6,7^, overexpression of microRNAs^8-13^, or delivery of modified RNAs has been shown to induce functional recovery in the infarcted heart in mice^14,15^. Although these gene transfer methods provide an important proof-of-concept for a new therapeutic approach to heart regeneration, due to concerns about safety and invasiveness, small molecule inducers for cardiomyocyte proliferation and cardiac regeneration have been sought eagerly^15-19^. However, despite much attention, pharmacological approaches to the induction of heart regeneration through activation of cardiomyocyte proliferation in adult mammals remain elusive.

Postnatal cardiomyocyte cell cycle arrest^6,20^, and subsequent hypertrophy as well as polyploidization, are regulated by several types of cyclin/CDKs^21-24^. Cyclin/CDKs also regulate the adult cardiomyocyte cell cycle^22,25^. Importantly, overexpression of CDK1, CDK4, cyclin B1, and cyclin D1 promotes cardiomyocyte proliferation in adult heart, and enhances the regeneration of the myocardium after myocardial infarction^26^. Thus, pharmacological activation of cyclin/CDKs potentially stimulates cardiomyocyte proliferation and heart regeneration.

## Results and Discussion

To induce cardiomyocyte proliferation by drug administration, we focused on benzyl isothiocyanate (BITC), which is known to regulate the cell cycle in a dose-dependent manner. In various types of cancer cells, BITC acts as an anti-proliferative and pro-apoptotic factor, while interestingly in normal cells the cytotoxic effects of BITC have not been observed^27-29^ and in some cases it acts as pro-proliferative factor^30^. The effects of BITC on normal cardiomyocyte proliferation have thus been unknown. First, we subcutaneously administered 20 mg/kg/day of BITC to neonatal mice from postnatal day 0 (P0) to P6, and examined the number of proliferating cardiomyocytes at P7. The ratio of heart weight per body weight was not changed, and cardiomyocyte size showed a trend toward a decrease in between BITC-treated mice and the control (Fig 1A). The double staining using cTnT and cell cycle markers suggests that cell proliferation was significantly promoted in cardiomyocytes (Fig 1B) but not in non-cardiomyocytes such as macrophages, endothelial cells, and fibroblasts (Figs EV1A and B, EV 2A and B) at P7 in the BITC-treated mice compared with control DMSO-treated ones. Interestingly, cardiomyocyte proliferation did not change upon treatment with 50 or 100 mg/kg/day of BITC (Fig EV1C), suggesting that BITC promotes cardiomyocyte proliferation in a dose-dependent manner. Next, in an attempt to identify the target of BITC, we examined cyclin-dependent kinases (CDKs) because a previous study revealed that cyclin and CDKs are important for cardiomyocyte proliferation^26^. Protein amount of Cyclin D1 and CDK 4 were not changed, while phosphorylated CDK1 (pCDK1) was increased in BITC-treated heart (Fig 1C). Based on this result, we examined whether BITC administration activates CDK1 *in vivo*. Immunohistochemical analysis revealed that pCDK1 in myocardial nuclei was increased in the BITC-administered heart at P7 (Fig 1C and D). This indicates that BITC induces the activation of CDK1, which may well contribute to cell cycle progression.

**Figure 1.**
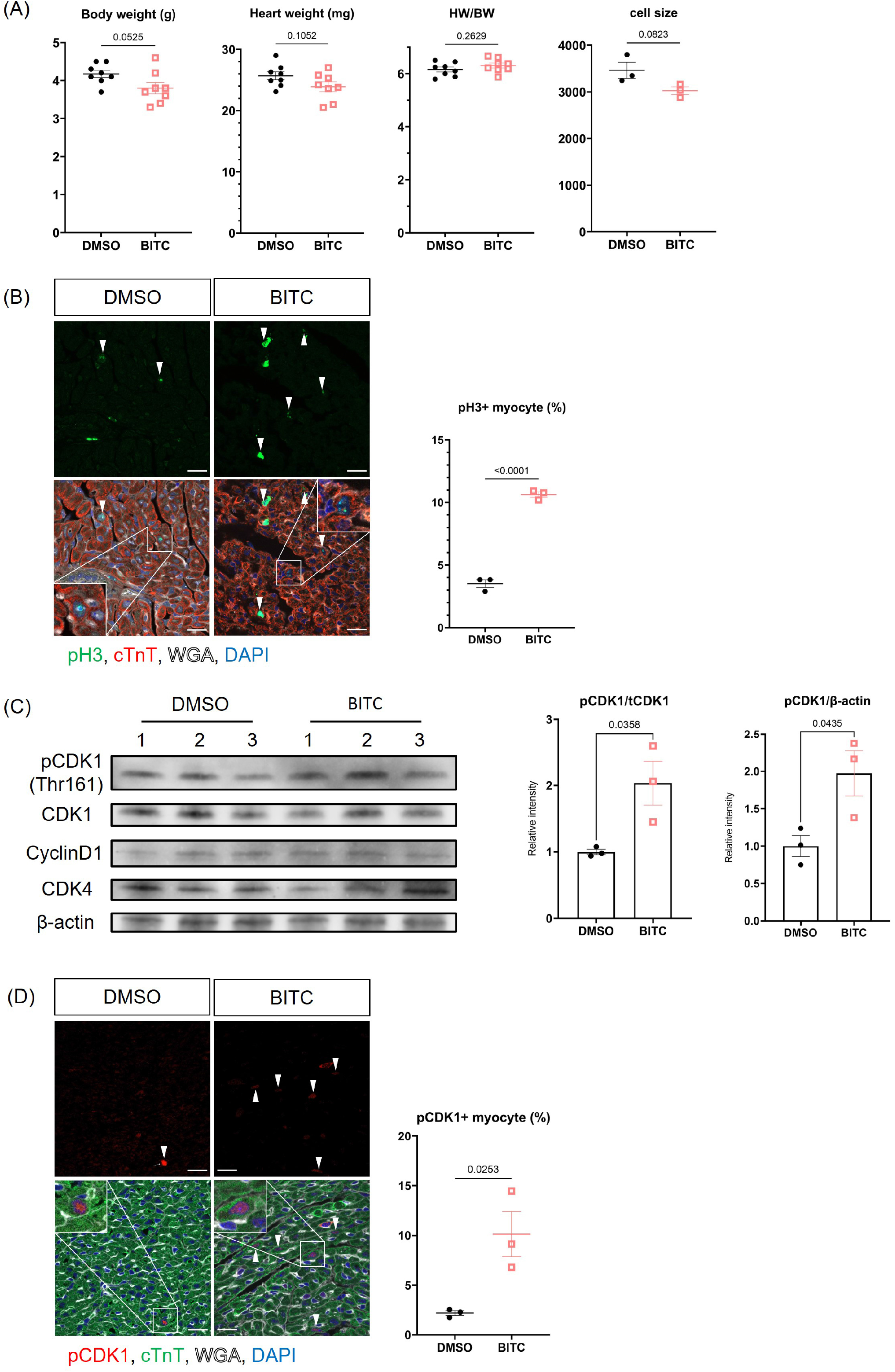
(A) Body weight, heart weight, heart weight normalized with body weight (HW/BW), and the cellular size of cardiomyocyte of DMSO or 20 mg/kg BITC administered mice at P7 (N=8 each for body weight and heart weight, N=3 each for the cellular size). (B) To quantify the number of pH3-positive cardiomyocytes, BITC-(20 mg/kg) or DMSO-administered heart sections were stained with pH3 (green) and cardiac troponin T (cTnT, red) antibodies (N=3 each). Arrowheads indicate pH3- and cTnT-double positive cells. (C) To quantify the amount of CDKs and cyclins in DMSO or 20 mg/kg BITC administered heart at P7, we performed Western blot analysis using anti-pCDK1, CDK1, CDK4, and cyclinD1 antibodies (N=3 each). (D) To quantify the number of pCDK1-positive cardiomyocyte nuclei, we used immunofluorescence of pCDK1 (red) and cTnT (green) in BITC- or DMSO-administered heart at P7 (N=3 each). Arrowheads indicate pCDK1- and cTnT-double positive cells.

To test whether BITC induces cardiomyocyte proliferation directly, we treated neonatal murine ventricular cardiomyocytes (NMVCs) with BITC. Interestingly, after treatment with 10-100μM BITC, only 50 μM of BITC showed a significant induction of cardiomyocyte proliferation compared with control (Fig 2A and B). Similarly, treatment with 50 μM of BITC on human iPS cell-derived cardiomyocytes (hiPSC) resulted in a tendency toward increased cell proliferation compared with control, although this enhanced proliferation was not statistically significant, most likely due to variability among hiPSC batches (Fig EV3). These results indicate that BITC directly induces cell proliferation in cardiomyocytes, rather than via an indirect systemic effect.

**Figure 2.**
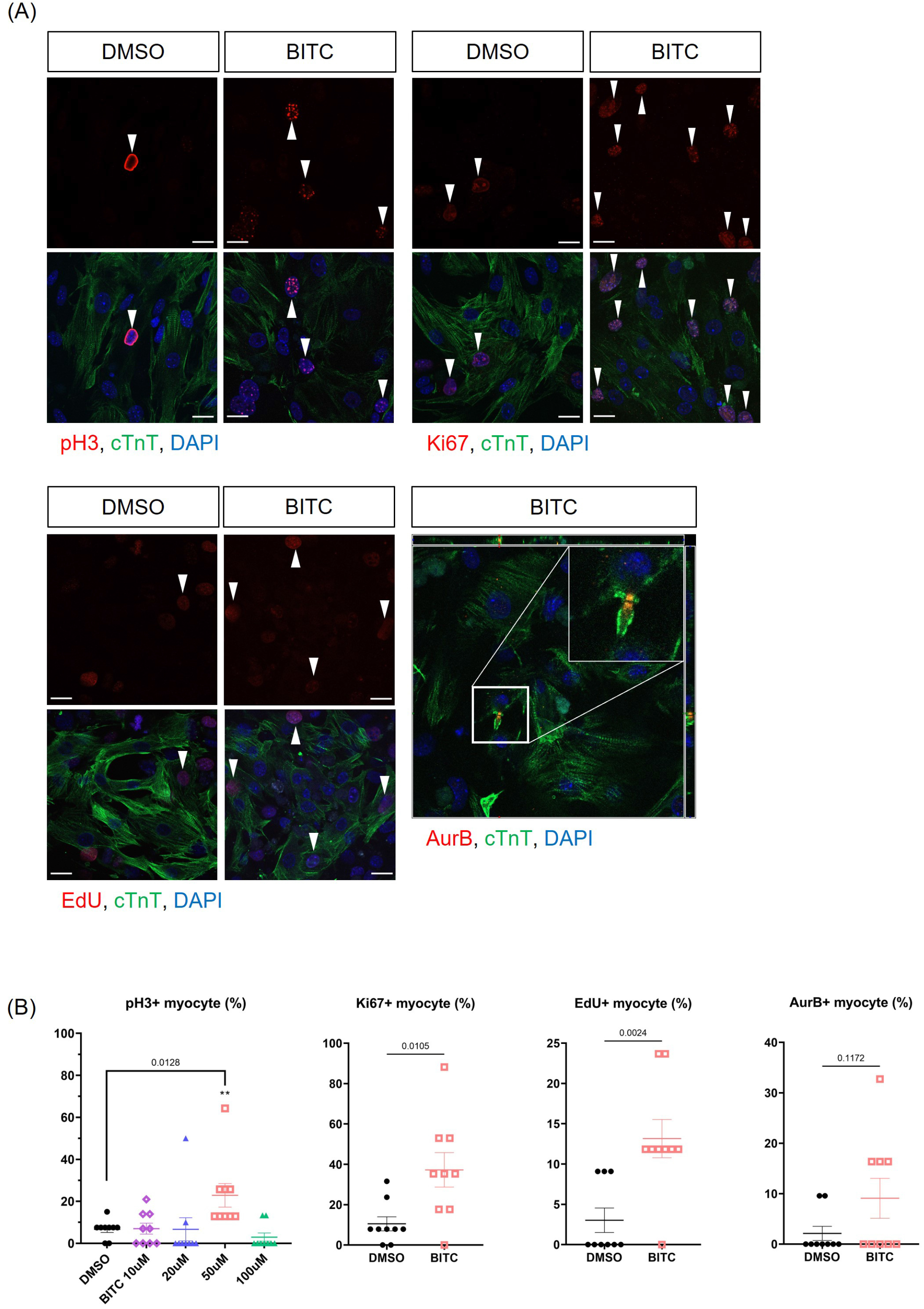
(A) Representative images of the primary culture of neonatal murine ventricular cardiomyocytes (NMVCs) immunostained with markers for cell cycle progression [pH3, Ki67, AurB, EdU (red) and cTnT (green)] upon DMSO or 50 μM BITC treatment. Arrowheads indicate cardiomyocytes positive for markers for cell cycle progression. (B) The ratio of pH3, Ki67, AurB, or EdU-positive cardiomyocytes was quantified (N=9 each).

Based on these results, we hypothesized that BITC treatment enables heart regeneration after myocardial infarction (MI) through enhanced cardiomyocyte proliferation. To test this hypothesis, we induced MI in P7 mice by permanent ligation of the left anterior descending coronary artery (LAD), and subsequently administered 20 mg/kg/day of BITC or control DMSO from P8 to P14 (Fig 3A). At P28, the ratio of heart weight per body weight was not changed between BITC-treated mice and the control mice (Fig EV4A), on the other hand, BITC-treated mice showed significantly recovered cardiac function (Figs 3B, EV3C) as well as an increased number of proliferating cardiomyocytes (Figs 3C, EV4B). Moreover, the fibrotic scar was significantly decreased in BITC-treated mice compared with DMSO-treated ones (Fig 3D and E), collectively suggesting that BITC administration induces cardiomyocyte proliferation, thereby allowing heart regeneration after MI in neonatal mice even after the regenerative period.

**Figure 3.**
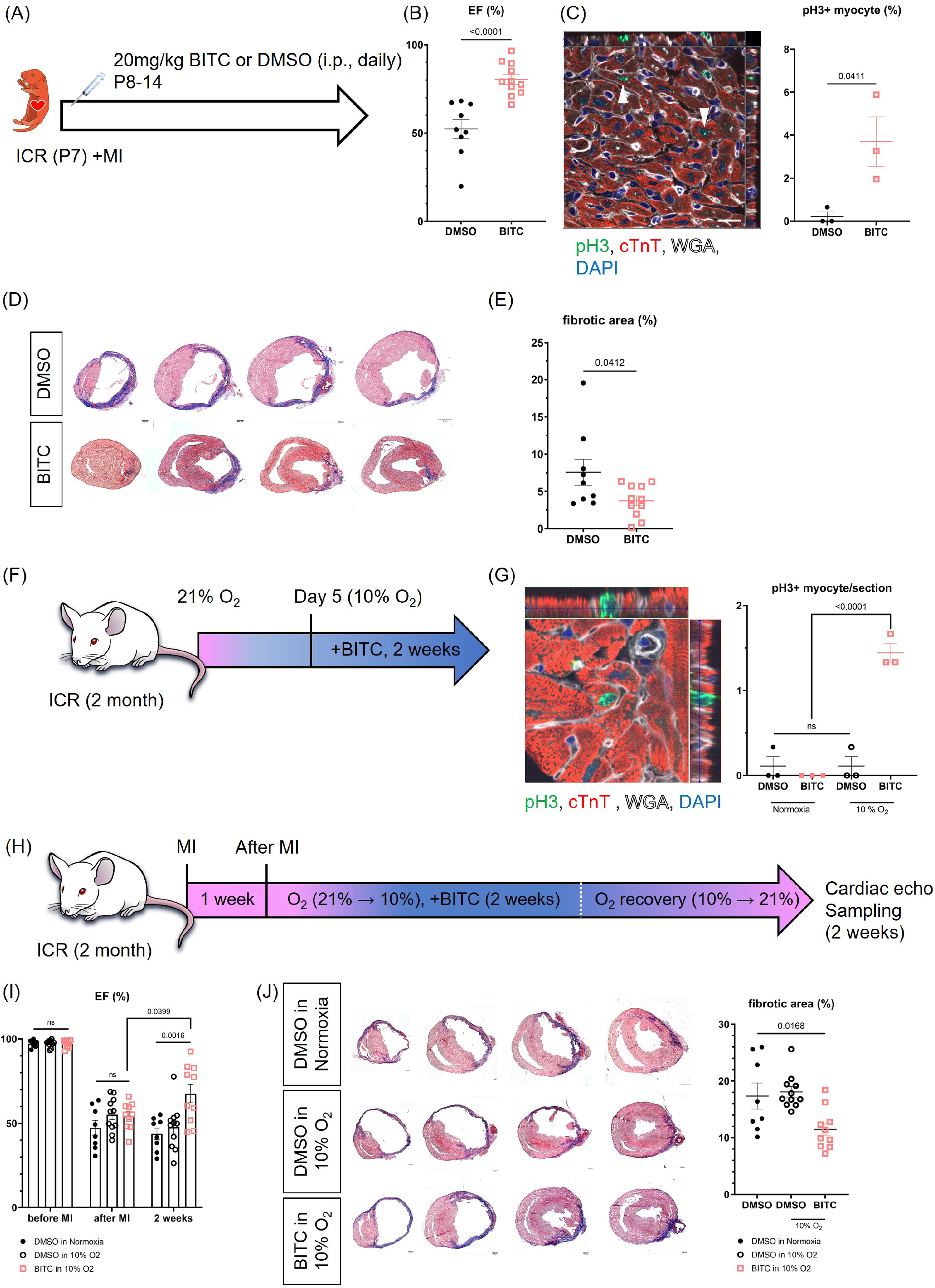
(A) Schematic of the experiment. After MI surgery at P7, BITC or DMSO was administered intraperitoneally once a day from P8 to P14. (B) Left ventricular ejection fraction (EF) was measured by echocardiography blindly at P28 (N=9 for DMSO, N=11 for BITC). (C) To quantify the number of pH3-positive cardiomyocytes, BITC-(20 mg/kg) or DMSO-administered MI heart sections were stained with pH3 (green) and cTnT (red) antibodies (N=3 each). Arrowheads indicate pH3- and cTnT-double positive cells. (D) Representative images of the histological sections of MI heart stained with Masson’s trichrome staining are shown. (E) The quantification of fibrotic area (N=9 for DMSO, N=11 for BITC) (F) Schematic of the experiment. Adult ICR mice were exposed to 10% O_2_ or normal conditions and injected with BITC or DMSO once a day for two weeks. (G) A representative image of pH3-positive cardiomyocytes taken with z-stack imaging with confocal microscopy and quantification of the number of pH3-positive cardiomyocytes (N=3 each). (H) Schematic of the experiment. Adult ICR mice were performed MI surgery, exposed to 10% O_2_ or normal conditions, and injected with BITC or DMSO once a day for two weeks. (I) Left ventricular ejection fraction (EF) was measured by echocardiography blindly after BITC administration. (J) Representative images of the histological sections of MI heart stained with Masson’s trichrome staining and the quantification of fibrotic area (N=8 for DMSO in normoxia, N=11 for DMSO in 10% O_2_, N=10 for BITC in 10% O_2_).

From a therapeutic standpoint, it is important to know whether cell cycle reentry is induced by BITC administration in the adult heart. In order to evaluate the effect of BITC treatment on adult cardiomyocytes, we administered 20 mg/kg/day of BITC intraperitoneally to 2-month-old ICR mice for 14 days. In parallel, we placed these mice in a 10% O_2_ environment based on findings from a previous study showing that a hypoxic environment aids cell cycle reentry in the adult heart (Fig 3F)^3^. pH3-positive cardiomyocytes were almost absent in the DMSO- or BITC-treated heart under the normal oxygen condition, nor were they present in the DMSO-injected heart in the 10% O_2_ condition in consistent with previous findings (Fig 3F and G)^3^. In contrast, pH3-positive cardiomyocytes were significantly increased after BITC administration in the hypoxic environment (Fig 3F and G), indicating that a combination of BITC administration and mild hypoxia exposure induces cell cycle reentry in the adult heart. To examine whether this enhanced cardiomyocyte proliferation contributes to heart regeneration in adult mice, we induced MI in 2-month-old mice and treated them with BITC in 10% O_2_ environment for 14 days, and then mice were recovered to a normoxic condition (Fig 3H). As a result, remarkably, cardiac function was significantly recovered (Figs 3I, EV5B) and the fibrotic area decreased (Fig 3J). No overt inflammation or tumorigenesis was observed upon treatment with BITC in combination with mild hypoxia (Fig EV5C). These results strongly indicate that BITC administration induces heart regeneration in adult mice.

Taken together, our data reveal a beneficial effect of BITC on heart regeneration. BITC treatment enhances cardiomyocyte proliferation through the activation of CDK1 in the neonatal heart. In addition, BITC administration in combination with mild hypoxia in adult mice is sufficient to induce cardiomyocyte cell cycle reentry, enabling regeneration of the infarcted area and recovery of cardiac function after MI. Given BITC administration induces the activation of CDK1 but not CDK4 or cyclin D1, it is suggested that BITC is not sufficient to induce cardiomyocyte proliferation, but rather requires partial metabolic reprogramming by mild hypoxia^22,31^. The present study suggests that pharmacological activation of the CDK pathway with BITC concurrently with the activation of hypoxia-related signaling pathways may enable researchers to establish a novel strategy to induce cardiac regeneration in patients with heart disease.

## Materials and Methods

### Mice

All protocols were approved by the Institutional Animal Care and Use Committee of RIKEN Center for Biosystems Dynamics Research. Wildtype ICR mice were purchased from CLEA Japan, Inc. and Oriental Yeast Co., Ltd. All experiments were performed on age-matched mice with approximately same number of males and females.

### Murine model of myocardial infarction

Induction of anterior wall myocardial infarction was undertaken using a protocol previously described^3^. Briefly, postnatal day (P) 7 mice were anesthetized with hypothermia on ice, and we ligated the left anterior descending coronary artery with polypropylene suture (Ethicon, EP8707H) after thoracotomy. Neonates were warmed on a hot plate (42 °C) after the surgery. Two-month-old mice were anesthetized with isoflurane, endotracheally intubated, and ventilated using a volume controlled ventilator during the operation of permanent LAD ligation with prolene suture (Ethicon, K890). Subsequently, the chest was closed using 5-0 silk sutures (Ethicon, J421), and the skin was closed using adhesive glue, Vetbond (3M, 1469SB). The mouse was then extubated and recovered on a hot plate.

### Echocardiography

Assessment of *in vivo* left ventricular dimensions and function in conscious mice were undertaken with Affiniti50 with an L15-7io transducer (Phillips) as previously described^3^.

### Hypoxia exposure

Exposure of adult mice to hypoxia was performed as described previously^3^. Briefly, mice were placed in a hypoxia chamber (COY laboratory) and the oxygen concentration was gradually decreased to 10% over 5 days. After 2 weeks of 10% O_2_ exposure in combination with drug administration, oxygen concentration was gradually recovered to 21%.

### Human iPS cell derived cardiomyocyte

The hiPSC lines GCaMP3-253G1^32,33^, was used in the present study. The maintenance of hiPSCs and differentiation of cardiovascular cells were conducted in accordance with previous studies^34-36^, with modifications. In brief, hiPSCs were expanded and maintained with StemFit AK02N medium (AJINOMOTO, Tokyo, Japan). At confluence, the cells were dissociated with TrypLE Select (Thermo Fisher Scientific, Waltham, MA, USA), dissolved in 0.5 mM ethylenediaminetetraacetic acid in PBS (1:1) and passaged as single cells (5,000 – 8,000 cells/cm^2^) every 5–7 days in AK02N containing iMatrix-511 silk (FUJIFILM Wako Pure Chemical Corp., Osaka, Japan) (0.125 µg/cm2) (uncoated laminin fragment) and ROCK inhibitor (Y-27632, 10 µM, FUJIFILM Wako)^37^. Penicillin–streptomycin (Thermo Fisher Scientific) (10,000 U/mL) (1:100 dilution) was used as required. For cardiovascular cell differentiation, single hiPSCs were seeded onto Matrigel-coated plates (1:60 dilution) at a density of 300,000–400,000 cells/cm^2^ in AK02N with Y-27632 (10 µM). At confluence, the cells were covered with Matrigel (1:60 dilution in AK02N) one day before induction. We replaced the AK02N medium with RPMI□+□B27 medium (RPMI 1640, Thermo Fisher; 2 mM L-glutamine, Thermo Fisher; 1×□B27 supplement without insulin, Thermo Fisher) supplemented with 100 ng/mL Activin A (R&D, Minneapolis, MN, USA) and 3–5 µM CHIR99021 (Tocris Bioscience, Bristol, UK) (differentiation day 0; d0) for 24 h, which was followed with supplementation with 10 ng/mL bone morphogenetic protein 4 (BMP4; R&D) and 10 ng/mL basic fibroblast growth factor (bFGF) (d1) for 4 days without a culture medium change. At d5, the culture medium was replaced with RPMI□+ □B27+insulin medium (RPMI 1640, Thermo Fisher; 2 mM L-glutamine, Thermo Fisher; 1×□B27 supplement with insulin, Thermo Fisher) supplemented with 2.5 µM IWP4 (Stemgent, Cambridge, MA, USA) and 5 µM XAV939 (Merck, Kenilworth, NJ, USA). The culture medium was refreshed with RPMI□+□B27+insulin medium every other day. Beating cells appeared at d11 to d15. Cardiomyocytes were purified with a specified medium [DMEM without glutamin, glucose and pyruvate, 4 mM L-lactic acid, and 0.1% BSA (Takara as described previously ^38^)]. The cells were fixed for immunostaining, and the images of cells were randomly recorded from nine different places. Cardiomyocyte differentiation, selection, and drug treatments were performed three times independently (N = 3).

### Primary culture

For cardiomyocyte isolation, 15 hearts were harvested from wildtype P1 ICR mice with approximately same number of males and females, and the ventricles were minced with iris scissors. The pieces of tissue were digested using the enzyme mix from a Neonatal Heart Dissociation Kit, mouse and rat (Miltenyi Biotec 130-098-373) at 37 °C with gentle pipetting. The suspension was added to 10% FBS (Thermo 10270106) in DMEM/F12 (Thermo 11320-033) and filtered through a 70-μm cell strainer (Falcon 352350), and then centrifuged (600*g*, 5 minutes at 25 °C). Precipitated cardiomyocytes were resuspended to 1 ml PBS (Wako 048-29805), treated with 10 ml Red Blood Cell Lysis Solution (Miltenyi 130-094-183) and centrifuged (600*g*, 5 minutes at 25 °C). The cell pellet was resuspended with culture medium [20% FBS, 5% horse serum (gibco 16050-130), 2 mM L-glutamine (gibco 25030-081), 0.1 mM NEAA (gibco 11140-050), 3 mM sodium pyruvate (gibco 11360-070), 1 μg bovine insulin (Sigma I0516) in DMEM/F12] and incubated on a non-coated dish to remove the adhesive cells (e.g. fibrotic cells at 37 °C for 90 minutes twice. We plated 4*10^5^ cells/well of isolated cardiomyocytes to fibronectin (cosmo bio SFN01)-coated 4 well chamber slides (Thermo Fisher 154526) and cultured them in a 5% CO_2_ incubator. We changed the culture medium 16 hours later to remove debris, and continued exchanging the medium once every other day during the culture period. The cells were fixed for immunostaining, and the images of cells were randomly recorded from nine different places (N = 9).

### Drug administration

Benzyl isothiocyanate (BITC SIGMA 252492) was dissolved in 10% dimethyl sulfoxide (DMSO WAKO 043-29355)/PBS (nacalai 27575-31), and 10-100 mg/kg/day of BITC was administered to neonatal mice by subcutaneous (s. c.) injection or adult mice by intraperitoneal (IP) injection. Neonatal murine ventricular cardiomyocytes in primary culture and human iPS cell-derived cardiomyocytes (hiPSC) were treated with 10-100 μM BITC (dissolved in 0.1% DMSO culture medium) for 6 hours.

### Immunostaining

Harvested hearts were fixed with 4% paraformaldehyde/PBS overnight at 4□°C, and embedded in tissue freezing medium (Genetal Data TFM-C) following replacement with 30% sucrose/PBS at 4□°C overnight, and cut to 8-μm thickness. For immunostaining, antigen retrieval was performed with epitope retrieval solution (IHC World IW-1100-1L) and a steamer (IHC-Tek Epitope Retrieval Streamer Set). Subsequently, tissue sections were blocked with 10% donkey serum/0.3 % TritonX-100/PBS, and incubated with primary antibodies (cTnT: Thermo Fisher MS295P, 1:100 or BD Pharmingen 564766, 1:500, CD31: abcam ab28364, 1:100, vimentin: abcam ab92547, 1:100, CD68 [FA-11]: Bio-Rad MCA1957, 1:200, pH3: Thermo Fisher 06-570 or SIGMA H9908, 1:100, Aurora B: Sigma A5102, 1:800, Ki67: Thermo Fisher 14-5698-82, 1:1000, pCDK1: abcam ab201008, 1:400) overnight at 4□°C, followed by incubation with corresponding secondary antibodies conjugated to Alexa Fluor 488 or 555 (Invitrogen A21202, A21206, A31570, and A31572), wheat germ agglutinin (50□mg/ml, Thermo Fisher W32466), and DAPI (nacalai, 11034-56) at room temperature. The slides were mounted in antifade mounting medium (nacalai, 12593-64).

### EdU labeling and detection

EdU (Wako 050-08844) was dissolved in PBS (50 mg/kg, *in vivo* IP injection) or culture medium (2 μM, *in vitro*), and stained with a Click-iT™ EdU Cell Proliferation Kit (Invitrogen C10338). Protocol is commercially described.

### Masson’s trichrome staining

TFM-embedded MI heart sections were cut to 10-μm thickness and stained with a Trichrome stain kit (ScyTek Laboratories TRM-1-IFU). Protocol is commercially described. The ratio of fibrotic scar area (Aniline Blue stained area per whole ventricular area) was quantified with ImageJ software from serial trichrome sections taken from the point of ligation all the way till the tip of the ventricular apex at intervals of 400 μm.

### Protein Extraction from Heart Tissue and Western Blotting

Freshly frozen hearts were homogenized in RIPA buffer containing phosphatase inhibitor cocktail I (Wako) and protease inhibitor tablet mini (Roche, cOmplete™, Mini), using a hand held homogenizer on ice for 1 min. Homogenates were then centrifuged at 12000 RPM for 20 min at 4°C, to remove insoluble material. Protein concentration was quantified using Pierce BCA protein assay kit (Pierce Biotechnology). Aliquots containing 5 µg protein were separated by 10% SDS-PAGE, transferred onto nitrocellulose membrane using Trans-Blot Turbo Transfer System (Bio-Rad) and Trans-Blot Turbo Mini 0.2 µm PVDF Transfer Packs (Bio-Rad). Membranes were blocked with casein- and phosphatase-free blocking solution (Blocking One-P, Nacalai Tesque) at room temperature for 20 min, and reacted with different primary antibodies at 4°C overnight. Subsequently, membranes were washed three times with TBS-T for 5 min each and then incubated with horseradish peroxidase (HRP)-conjugated secondary antibodies (anti-mouse/rabbit) (MBL, 330, 458) for 1 hour at room temperature. The primary antibodies used for western blotting are as follows: Thr^161^-phosphorylated CDK1 (Abcam, ab201008, EPR19546; 1:1000), CDK1 (Abcam, ab32094, YE324; 1:1000), CyclinD1 (Abcam, ab134175, EPR2241, 1:100000), CDK4 (Abcam, ab199728, EPR17525; 1:2000), β-actin (Proteintech, 66009-1-lg, 2D4H5; 1:1000). The immunoreactive protein bands were visualized using Chemi-Lumi One Super reagent (Nacalai Tesque) and imaged using an LAS-4000 luminescent image analyzer (FUJIFILM). Quantification analysis of the immunoreactive protein band signal intensity was performed with Fiji/ImageJ software.

### Microscopy

Images of immunohistochemical, histological, and immunocytochemical samples were taken with Olympus BX53 and Carl Zeiss LSM800 confocal microscope.

### Data analyses

Data are means ± SEM, the *P* values were determined with student’s t test, and one-way ANOVA or two-way ANOVA were applied for multiple comparison. All scale bars represent 20 μm. No statistical method was used to estimate sample size.

## Acknowledgement

We thank RIKEN BDR Laboratory for Animal Resources and Genetic Engineering for animal husbandry, and Y. Shiba for providing a human iPS cell line.

## Authors contributions

A.S. designed and performed the experiments, analyzed the data, and wrote the manuscript. M.K. and K.M. performed human iPS cell-related experiments. Y.S. performed studies related to western blot. H.M. performed and supervised human iPS cell-related experiments. W.K. conceived the study, designed the experiments, analyzed the data and contributed to manuscript preparation.

## Figure legends

**Expanded View Figure 1.**
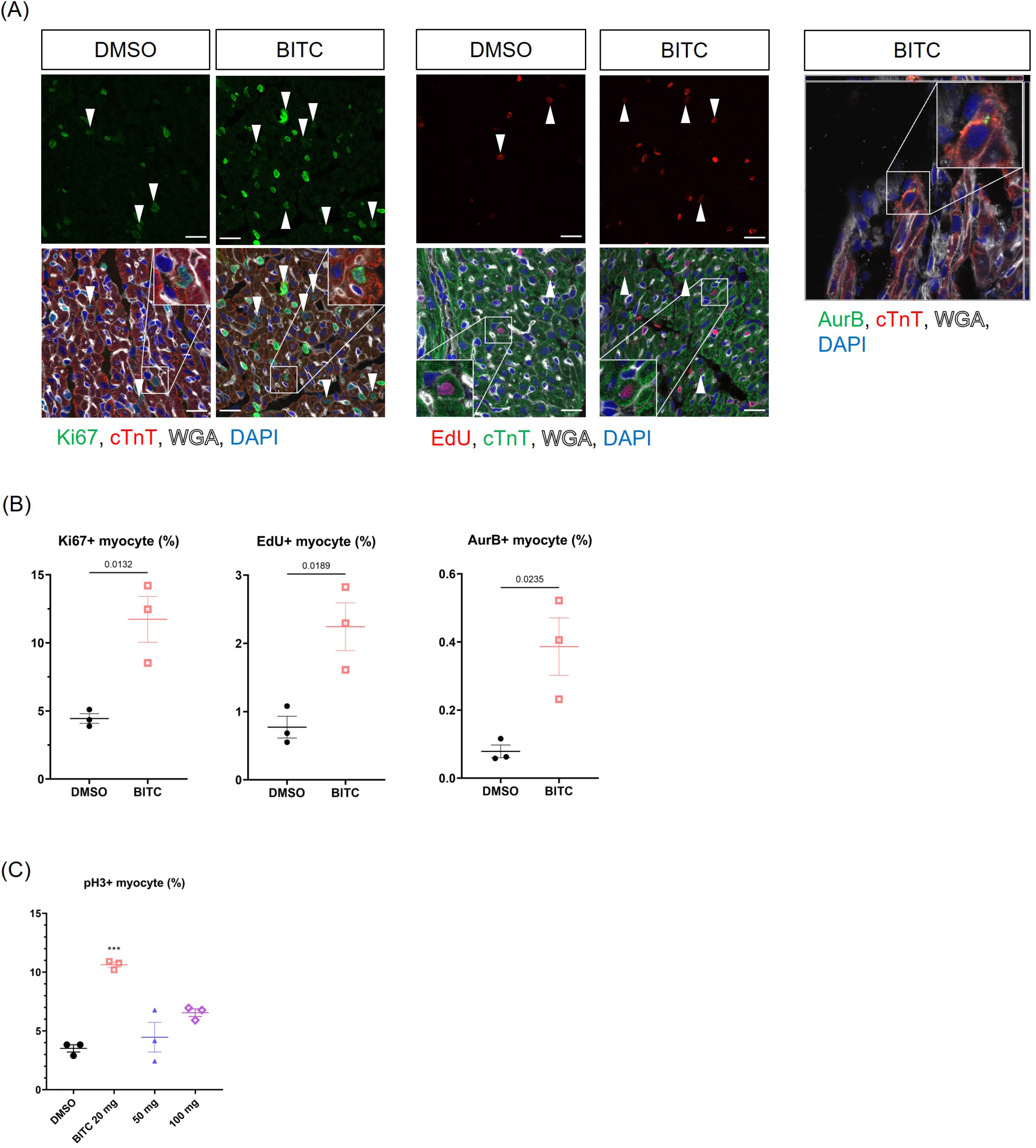
(A) Representative images of cell cycle marker (Ki67, AurB, or EdU)-positive cardiomyocytes. Arrowheads indicate cardiomyocytes positive for cell cycle markers. (B) The quantification of cell cycle marker positive cardiomyocytes from (A) (N=3 each).

**Expanded View Figure 2.**
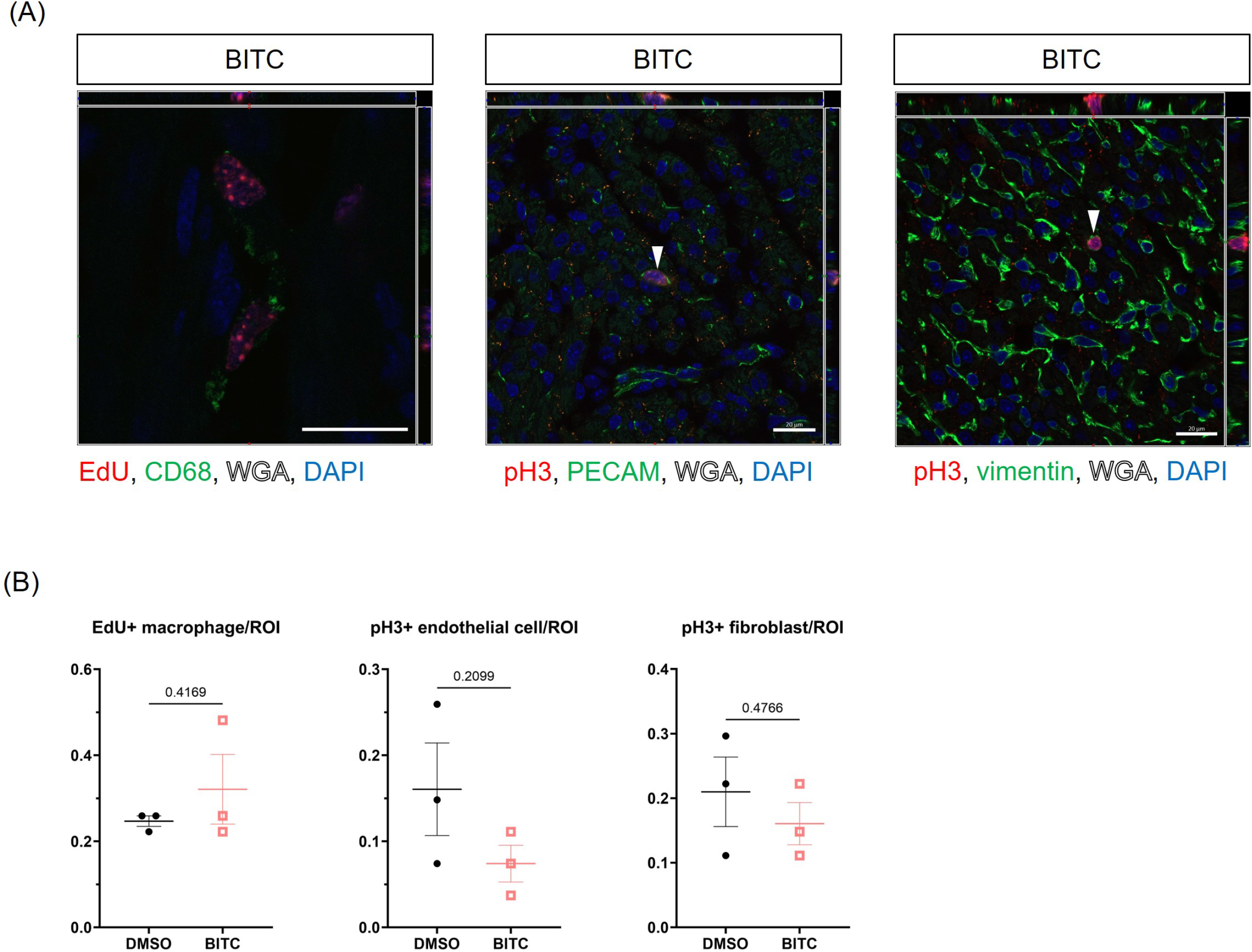
(A) Representative images of co-staining of cell cycle markers and markers for macrophages (CD68), vascular endothelial cells (PECAM), and fibroblasts (vimentin) in BITC- (20 mg/kg) administered heart sections. Arrowheads indicate non-cardiomyocytes positive for cell cycle markers. (B) The quantification of cell cycle marker positive macrophages, vascular endothelial cells, and fibroblasts in BITC- (20 mg/kg) or DMSO-administered heart sections (N=3 each).

**Expanded View Figure 3.**
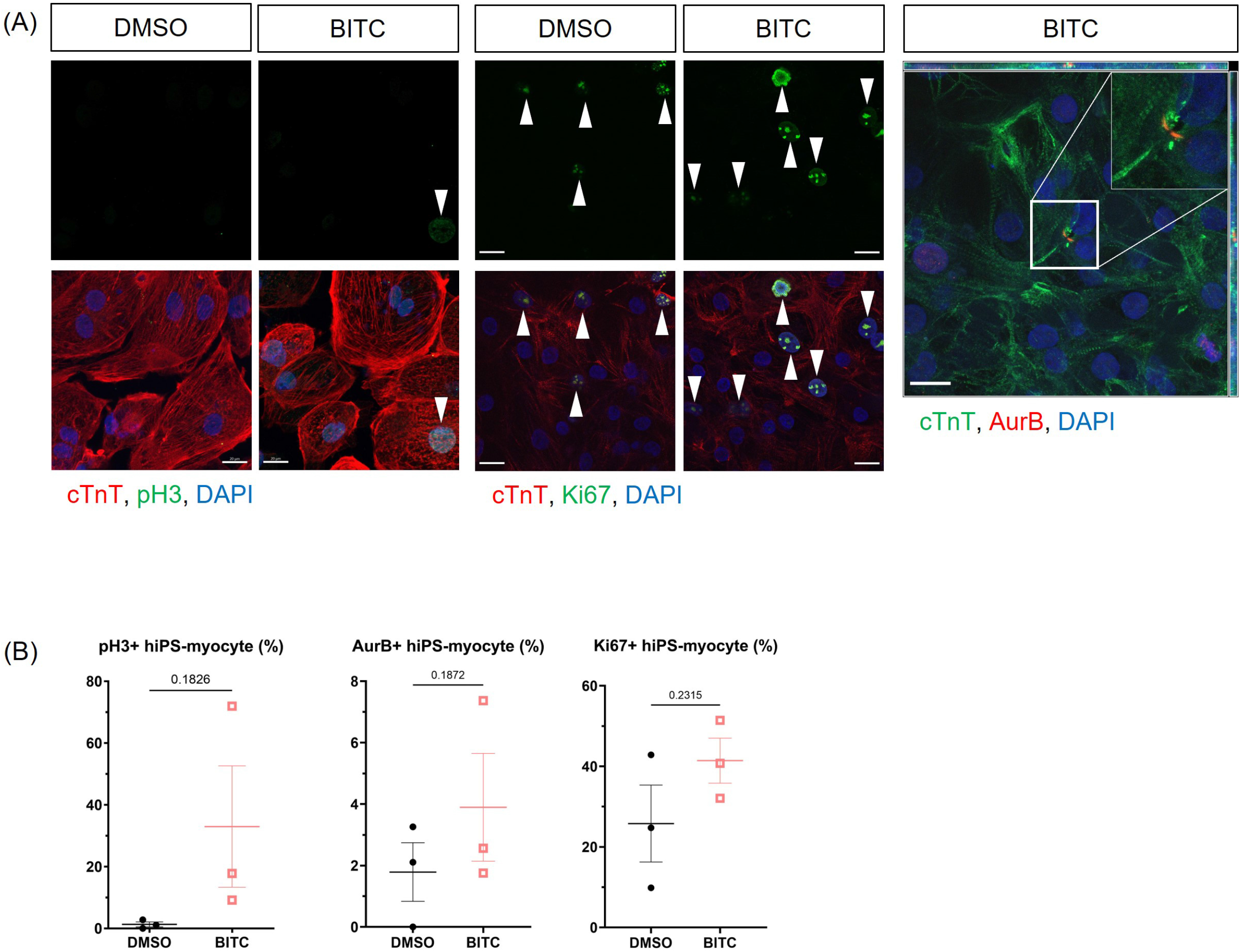
(A) Representative images of hiPSC-CMs treated with 50 μM BITC or DMSO. Arrowheads indicate hiPSC-CMs positive for cell cycle markers. (B) The quantification of cell cycle marker positive cardiomyocytes from (A) (N=3 each).

**Expanded View Figure 4.**
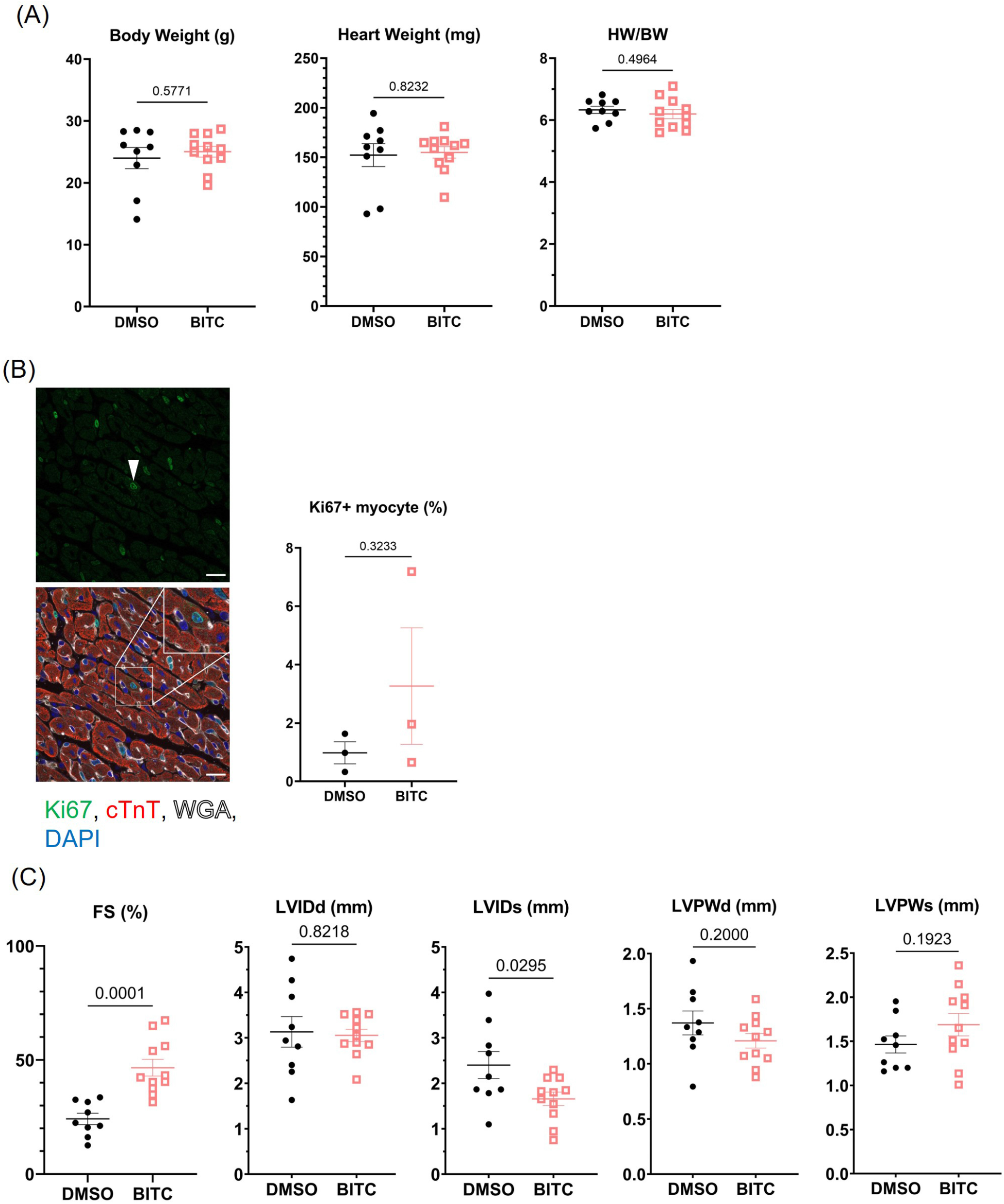
(A) Body weight, heart weight, and HW/BW ratio of DMSO or 20 mg/kg BITC administered MI mice at P28 (N=9 for DMSO, N=11 for BITC). (B) To quantify the number of Ki67-positive cardiomyocytes, BITC- or DMSO-administered MI heart sections were stained with Ki67 (green) and cardiac troponin T (cTnT, red) antibodies (N=3 each). Arrowheads indicate Ki67-positive cardiomyocytes. (C) Fraction shortening (FS), left ventricular internal diameter (LVIDd, LVIDs), and left ventricular posterior wall thickness (LVPWd, LVPWs) of P7-MI heart were measured by echocardiography blindly at P28 (N=9 for DMSO, N=11 for BITC).

**Expanded View Figure 5.**
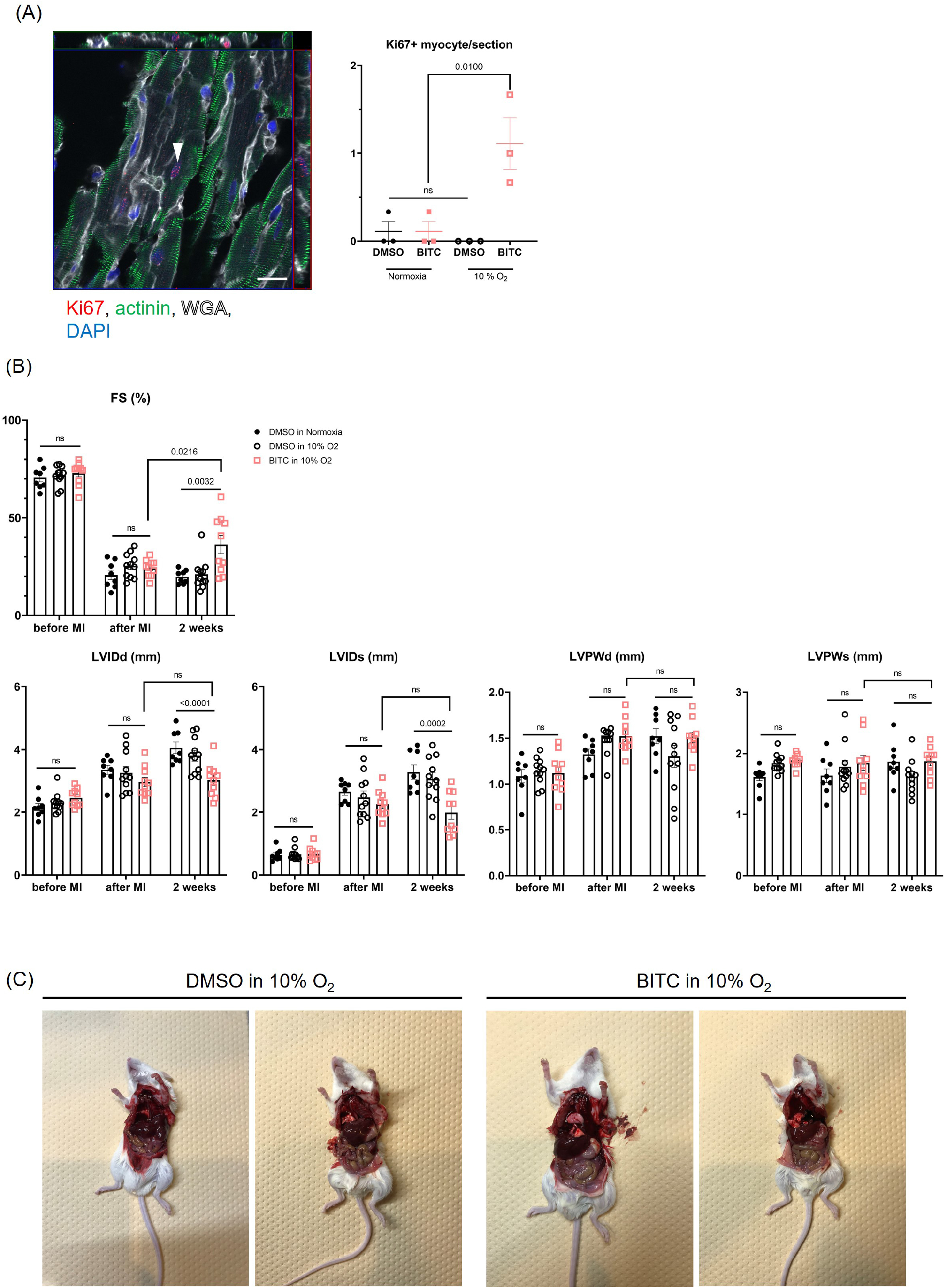
(A) A representative image taken with z-stack imaging with confocal microscopy and quantification of Ki67-positive cardiomyocytes (N=3 each). (B) Fraction shortening (FS), left ventricular internal diameter (LVIDd, LVIDs), and left ventricular posterior wall thickness (LVPWd, LVPWs) of adult MI heart were measured by echocardiography blindly (N=8 for DMSO in normoxia, N=11 for DMSO in 10% O_2_, N=10 for BITC in 10% O_2_). (C) Representative images of mice after DMSO/BITC administration and mild hypoxia exposure.

